# PEDF and Derived Peptides Prevent Apoptosis and Promote Differentiation of Retinal Photoreceptors

**DOI:** 10.1101/2021.01.01.425044

**Authors:** Germán Michelis, Olga Lorena German, Rafael Villasmil, Nora P. Rotstein, Luis Politi, S Patricia Becerra

**Author notes:** Corresponding authors: Luis Politi, Instituto de Investigaciones Bioquímicas de Bahía Blanca (INIBIBB) Florida 7000, 8000 Bahía Blanca Argentina, S. Patricia Becerra NIH-NEI, Building 6, Room 134, 6 Center Drive MSC 0608, Bethesda, MD 20892 USA.

## Abstract

Pigment epithelium-derived factor (PEDF) is a cytoprotective protein for the retina. We hypothesize that this protein acts on neuronal survival and differentiation of photoreceptor cells in culture. The purpose of the present study was to evaluate the neurotrophic effects of PEDF and its fragments in an *in vitro* model of cultured primary retinal neurons that die spontaneously in the absence of trophic factors. Results show that PEDF protected photoreceptor precursors from apoptosis, preserved mitochondrial function and promoted polarization of opsin enhancing their developmental process, as well as induced neurite outgrowth in amacrine neurons. These effects were abolished by an inhibitor of the PEDF receptor or receptor-derived peptides that block ligand/receptor interactions. While all the activities were specifically conferred by short peptide fragments (17 amino acid residues) derived from the PEDF neurotrophic domain, no effects were triggered by peptides from the PEDF antiangiogenic region. The observed effects on retinal neurons imply a specific activation of the PEDF receptor by a small neurotrophic region of PEDF. Our findings support the neurotrophic PEDF peptides as neuronal guardians for the retina, highlighting their potential as promoters of retinal differentiation, and inhibitors of retina cell death and its blinding consequences.

## Introduction

The importance of pigment epithelium-derived factor (PEDF), as both a neurotrophic factor and an antiangiogenic agent, for the retina has been well-documented *(Barnstable and Tombram-Tink, 2004; Bouck, 2002; Becerra and Notario 2013; Polato and Becerra 2018)*. Moreover, its deficiency is associated with progression of retinodegenerative diseases, such as retinitis pigmentosa, age-related macular degeneration and diabetic retinopathy *(Holenkmap et al., 2002; Ogata et al., 2004; Zhang et al., 2006)*.

PEDF is a glycoprotein and a member of the superfamily of serine protease inhibitors (SERPIN) that belongs to the subgroup of ‘non-inhibitory’ serpins (Steele et al., 1993; Becerra et al 1995). The monolayer of cells of the retinal pigment epithelium (RPE) secretes PEDF in an apicolateral fashion to be deposited in the interphotoreceptor matrix where it acts on survival of PHRs and is responsible for its avascularity (*Tombran-Tink et al., 1995; Becerra et al., 2004; Dawson et al., 1999; Cai et al., 2006; Michalcyzk et al., 2018*). We have mapped the regions that confer the neurotrophic and antiangiogenic activities to the multifunctional PEDF polypeptide. The region spanning between amino acid residue positions 44-77 of the human sequence, termed 34-mer, contains the antiangiogenic domain, while a region spanning between positions 78-121, 44-mer, contains the neurotrophic domain (*Becerra 2006*). Furthermore, a smaller synthetic mimotope, the 17-mer peptide (98-115 positions), designed from the 44-mer peptide, recapitulates the neurotrophic properties of the full-length PEDF protein of about 398 residues (*Kenealey et al., 2015*). The distinct 34-mer peptide is an antiangiogenic mimotope of PEDF devoid of neurotrophic activity (*Amaral and Becerra, 2010; Becerra et al 1995*). PEDF elicits cytoprotective effects by binding and activating its receptor PEDF-R *(Notari et al., 2006)*. Retinal cells express the gene for PEDF-R termed patatin-like phospholipase-2 (*Pnpla2*) (*Notari et al., 2006*). The pattern of *Pnpla2* expression in the mouse retina by laser capture microdissection reveals that the different layers of the retina express this gene (*Dixit et al, 2020*). In the albino rat retina PEDF-R distributes at higher levels in the inner segments of PHRs than in the rest of the tissue (*Notari et al., 2006; Subramanian et al., 2010*). The PEDF-R protein, also known as calcium-independent phospholipase A2 (iPLA2), is an enzyme with phospholipase A2 activity that liberates fatty acids and lyso-phospholipids from phospholipids (*Notari et al 2006; Jenkins et al., 2004*). Specifically, it hydrolyzes the sn-2 acyl bond of phospholipids thus releasing polyunsaturated fatty acids, preferentially docosahexaenoic acid (DHA, 22:6n-3) (*Pham et al., 2017*). While PEDF binding stimulates the PEDF-R enzymatic activity (*Notari et al 2006*), its inhibition by atglistatin abolishes the PEDF survival effects in retinal R28 cells (*Kenealey et al. 2015*). Interestingly, the phospholipase A2 inhibitor bromoenol lactone (*BEL*) inhibits PEDF-R activity and blocks both the PEDF-and DHA-mediated survival effects on retina R28 cells and PHRs undergoing serum-starvation-induced death and oxidative-stress-induced apoptosis, respectively (*Subramanian et al. 2013; German et al., 2013*). These similarities imply a direct action of PEDF on PHRs, likely via stimulation of PEDF-R to generate DHA as its neurotrophic lipid mediator. However, the direct effects of PEDF on PHRs and its mechanism of action on PHR development are unknown.

Most of the research to evaluate the effects of PEDF on PHRs has been performed in animal models of retinal degeneration *in vivo* (*Polato and Becerra, 2018*). However, the complexity of the interactions that occur in the native retina makes it difficult to evaluate the subcellular changes involved in biochemical pathways triggering the PEDF effects on PHRs, which remain to be elucidated. Purified neuronal retinal cultures are ideal to perform these types of studies. We have previously established primary cultures of purified neurons enriched in PHRs and amacrine neurons cultured in a complete chemically defined medium *(Politi et al., 1988)*. As it occurs *in vivo*, in the absence of their specific trophic factors, PHRs in these cultures undergo a developmental programmed cell death *(Politi et al., 1988). C*onversely, addition of growth factors, such as glial-derived factor (GDNF), fibroblast growth factor (FGF) *(Fontaine et al., 1998)*, or the lipid molecule docosahexaenoic acid (DHA, 22:6n-3) reverses this process to prevent death of PHRs in culture (*Politi et al 2001*). In addition to protecting PHRs against death by apoptosis, DHA promotes their differentiation in these cultures *(Rotstein et al., 1997; 1998*).

However, the neurotrophic potential of PEDF has not yet been tested in this primary cell culture system.

Using neuronal retinal cell cultures, our primary outcome was to examine the effects of PEDF on PHR development and survival. Here, we provide evidence that PEDF and its 17-mer, and 44-mer derived peptides protect PHRs, promote their differentiation and stimulate axonal outgrowth, selectively in amacrine cells. We discuss how the findings provide a more comprehensive view of the intracellular molecular events elicited by PEDF and its derived peptides on isolated PHRs and amacrine neurons in vitro.

## Materials and Methods

### Protein, peptides, and reagents

Recombinant human PEDF was produced and purified as outlined previously (*Stratikos et al 1996*). Human PEDF-R peptide P1 and human PEDF peptides 34-mer, 44-mer, and 17-mer were synthesized as described (*Kenealey et al., 2015*) (see Table S1). Dulbecco’s Modified Eagle Medium (DMEM) (31600034) (2015-2020), Hanks’s Balanced Saline Solution (HBSS) (14170112) (2015-2018), trypsin (15090046) (2015-2018), gentamicin (15750060) (2015-2017), distilled water (15230147) (2017-2020) and Penicillin-Streptomycin (15140122) (2015-2018) were from Gibco (US). Poly-ornithine (P3655) (2015, 2017, 2020), insulin (I4011) (2015, 2016, 2017, 2020), trypsin inhibitor (T6522) (2015-2018), hydrocortisone (H0888) (2015-2018), putrescine (P5780) (2015-2018), apotransferrine (T1147) (2015-2018), CDP-coline (90756) (2015-2018), CDP-ethanolamine (C0256) (2015-2019), progesterone (P8783) (2015-2018) were from Sigma (US).

### Animals

One-day-old albino Wistar rats (RRID:RGD_13508588) that were either bred in the animal facilities of INIBIBB and NIH or purchased through Charles River (Frederick, US) were used in all the experiments without distinction of sex, without any exclusion criteria set. Euthanasia was performed by decapitation. All proceedings concerning animal use were done in accordance with the guidelines published in the Guide for the Care and Use of Laboratory Animals. All animals were housed and handled in accordance with the ARVO statement for the Use of Animals in Ophthalmic and Vision Research, USA guidelines, and following the protocols approved by the Institutional Committee for the Care of Laboratory Animals from the Universidad Nacional del Sur (Argentina) (Protocols CICUAE-UNS: 058/2015, 059/2015, and 060/2015), as well as the National Eye Institute Animal Care and Use Committee (NEI-ASP 682).

No randomization was performed to allocate subjects in this study. For the in vitro experiments a total of 60 rat pups were used.

### Neuronal Cell Cultures

Culture dishes and wells were pretreated with poly-ornithine diluted in borate buffer (0.1 mg/mL) for at least an hour, and then with 25% Schwannoma conditioned medium overnight. Pure neuronal retinal cultures were obtained according to previously established procedures (*Politi et al., 1988; Rotstein et al, 1996*). Briefly, retinas were extracted from 1-day old pups, dissected and dissociated under mechanical and trypsin digestion. Cells were then re-suspended in a chemically defined medium and seeded on 35 mm- or 60 mm-diameter dishes (Greiner Bio-One 627160 and 628160) (2015-2019), according to the experimental design. Cultures were incubated at 36.5°C in a humidified atmosphere of 5% CO_2_. These studies were not pre-registered

### Immunocytochemical Assays

At day 5 after seeding, the cultures were fixed with 2% paraformaldehyde (Sigma, P6148) (2015-2019) for 30 min at room temperature, permeabilized with Triton X-100 0.2% for 10 min, incubated with primary antibodies for 1 h, followed by secondary antibodies conjugated to either Cy2, Cy3 (Jackson Immunoresearch, US) (2015-2017), Alexa 488 or 555 (Invitrogen, US) (2017-2019), diluted in PBS (All of the antibody combinations and concentrations can be found in Table 1 in the appendix). Nuclei were visualized with either Hoescht 33342 (Invitrogen, H3570) (2017, 2018) (20 µM); 4,6-diamidinole-2-phenylindole (DAPI) (Sigma, D9542) (2015-2017, 2019, 2020) (35 µM) or TOPRO-3 (Sigma, T3605) (2016) (2µM), incubated 1:1000 for 1 h. Between permeabilization and antibody incubations, samples were washed three times with PBS for at least 5 min each time. Unless specified, all incubations were done at room temperature. Any deviation from this protocol is indicated in their individual sections below.

**Table 1:** Immunocytochemistry data. Antibody combinations used in this paper, along with concentration and incubation time.

| Primary antibody* |  |  |  | Secondary antibody** |  |  |
| --- | --- | --- | --- | --- | --- | --- |
| Name | Company | Cat. Number | Dilution | Name | Company | Cat. Number |
| HPC-1 | Abcam | ab3265 | 1/40 | Goat anti-mouse (Cy2) | Invitrogen | 115-225-146 |
| PEDF-R (1) | Sigma-Millipore | ABD66 | 1/100 | Goat anti-rabbit (Alexa 555) | Invitrogen | A27039 |
| PEDF-R (2) | Cayman | 10006409 | 1/100 | Goat anti-rabbit (Alexa 555) | Invitrogen | A27039 |
| PEDF-R (3) | R&D | AF5365 | 1/100 | Goat anti-mouse (Alexa 488) | Invitrogen | A28175 |
| Rhodopsin | Sigma-Millipore | MAB5356 | 1/100 | Goat anti-mouse (Alexa 488) | Invitrogen | A28175 |
| Rhodopsin | Gift of R. Molday | N/A | 1/100 | Goat anti-mouse (Cy2) | Jackson Immunoresearch | 115-225-146 |
| Tuj1 | Abcam | ab24610 | 1/300 | Goat anti-mouse (Cy2) | Jackson Immunoresearch | 115-225-146 |
| Acetylated Tubulin | Abcam | ab18207 | 1/300 | Goat anti-rabbit (Cy3) | Jackson Immunoresearch | 115-165-003 |
| Na/K pump | Sigma-Millipore | 06-520 | 1/100 | Goat anti-rabbit/mouse (Alexa 555/488) | Invitrogen | A27039/A28175 |

\*Primary antibodies were diluted in PBS and the incubation time and temperature were 1 h at 25°C or 16 h at 4°C
\*\*Secondary antibodies were diluted 1:200 in PBS and the incubation time was 1 h at room temperature

### Microscopy

Images were acquired using laser confocal scanning microscopy (DMIRE2/TSCSP2 microscope; Leica, Wetzlar, Germany) with a 63X water objective, and processed (LCS software; Leica, 2006) or by the confocal microscopes (Olympus FV1000, Zeiss 700 and Zeiss 880, 2009). The images were analyzed and processed using the ImageJ software program and ZEN, the proprietary software of Zeiss. For quantification purposes, each experiment was performed at least three times, with each having three biological replicates (three dishes) per condition, and from which 10 random fields were photographed and analyzed.

### Cell identification

Neuronal cell types were identified by their morphology using phase contrast microscopy and by immunocytochemistry. Amacrine neurons were identified by immunocytochemistry with the monoclonal Anti-Syntaxin antibody. Under the microscope, we identified individual cell types as follows: 1) PHRs have a small (5-10 µm) round and dark cell body bearing a single neurite at one end, which usually ends in a conspicuous synaptic “spherule”, and a cilium at the other end, with or without an outer segment-like process; 2) Amacrine neurons have either single or multiple neurites, a bigger cell body (around 10-30 µm) and a broad morphological heterogeneity. The remaining neuronal cell types in culture represented less than 1% of total cells.

### Addition of PEDF and derived peptides

PEDF and its derived peptides, 17-mer, 44-mer and 34-mer were diluted in HBSS and aliquots were added to the cultures at 10 nM final concentration 48 h after seeding the cells. The 17-mer and 44-mer peptides from the neurotrophic domain were the effectors and the 34-mer peptide from the antiangiogenic domain of PEDF was used as a negative control. For suppressing the effects of PEDF and its derived peptides, a fragment of the ligand-binding site of PEDF-R, the P1 blocking peptide, diluted in HBSS was added at a P1-PEDF molar ratio of 10:1. Alternatively, a selective inhibitor of PEDF-R, Atglistatin (Sigma, SML1075) (2017-2020) at a 3.5 µM concentration (aliquoted and diluted in DMSO) was used. Both blocking peptide and inhibitor were preincubated in the cultures for 1 h before adding the effectors.

### Immunolabeling of PEDF-R

Cells were cultured in 4-chamber slides (Thermo Fisher Scientific, 154526) (2018). The fixed and permeabilized cells were incubated in a solution containing antibodies for PEDF-R for 1 h. For membrane marker detection, cells were stained with Wheat Germ Agglutinin (WGA) coupled to the fluorophore Alexa 555 (Thermo Fisher Scientific, W32464, US) (2018) for 10 min after fixation and before permeabilization and following the manufacturer’s instructions. Then the images of cells were captured with the microscopes Zeiss 700 and 880. Z-stacks images were acquired with a 0.6 micrometer step size. The resulting stacks were then processed with the ZEN software (V16.0.0.0).

### Evaluation of neuronal cell death

Cell death was determined by incubating the cultures with 7.5 µg/ml of Propidium Iodide (PI) (Sigma, P4170) (2016, 2017), diluted in the media for 30 min at 37°C. Then the cultures were washed twice with PBS, fixed, permeabilized, counterstained either with DAPI or TOPRO-3, and analyzed by using phase-contrast, or laser scanning confocal microscopy, as described above. PI-labeled cells were considered non-viable.

Apoptosis was determined with the ApopTag® Fluorescein In Situ Apoptosis Detection Kit (Sigma, S7110) (2017-2019), following the manufacturer’s instructions, and nuclei were counterstained with DAPI (diluted 1:100), incubated for 15 min.

To evaluate cell death by Flow Cytometry, we used APO-BrdU™ TUNEL Assay Kit (Invitrogen, A23210) (2017), following manufacturer’s instructions. These samples, along with positive and negative controls provided with the kit were analyzed in an Cytoflex NUV LX Flow Cytometer (Beckmann-Coulter) (2017). Additionally, we used Annexin V-Mitotracker kit (Thermofisher, V35116) (2017) for flow cytometry, following the manufacturer’s instructions. In both cases, DAPI was used to counterstain dead cells. Flow Cytometry data was analysed with CytExpert (Beckmann-Coulter) (V2.3) and FlowJo (BD Biosciences) (V10.0.4) software. Thresholds between quadrants were set by using the positive and negative control samples provided in the kit.

### Mitochondrial assays

Mitochondrial membrane potential was assessed by MitoTracker Red CMXRos (Invitrogen, M7512) (2016, 2017, 2019) prepared and used according to the manufacturer’s instructions. Briefly, Mitotracker was added at a 200 nM concentration to the cell cultures and incubated for 30 min. The cultures were then washed with PBS, fixed, permeabilized and their nuclei were counterstained with DAPI as described above, and cell labeling was then analyzed with a fluorescence microscope (Nikon Eclipse 2000, 2006). Cells with intense and discreet signal located on the axon hillock were selected as healthy.

### Rhodopsin determination

Rhodopsin expression in PHRs was assessed by immunocytochemistry by using the Rho4d1 and Rho4d2 antibodies (MAB5356 from Sigma-Millipore, and a generous gift of Dr. Robert Molday, respectively) (2016, 2017, 2019). Both showed equivalent staining patterns.

### Neurite outgrowth

To determine neurite outgrowth, pretreatment of dishes with the RN22 conditioned medium, which promotes neurite outgrowth, was omitted. The cells were fixed, permeabilized as described above and incubated with anti-Acetylated Tubulin or Tuj1 for 1 h. The length of the longest neurite and diameter of each cell were measured using ImageJ. The length was then divided by the diameter of its respective cell. A threshold was set, only considering the measurements belonging to cells whose longest neurite had a length of 4 or more cell diameters. Then ratios were plotted using GraphPad Prism (Version 8.2.0).

### Western Blotting

Cell cultures were lysed with RIPA buffer (Thermo Fisher Scientific, 89900) according to the manufacturer’s instructions, and then the lysates were kept frozen at −80°C until use. Protein concentration in the lysates was quantified by the Bradford assay (Invitrogen, 23225, US). An equal amount of protein for each lysate was loaded into NuPAGE 10% Bis-Tris gels (Thermo Fisher Scientific, NP0301, US) (2017-2020). The electrophoretic run was performed for 90 min at 125 volts with MOPS as the running buffer (Thermo Fisher Scientific, NP0001, US) (2017, 2018). The proteins in the gel were then transferred to a membrane using the iBlot 2 membrane transfer device (Invitrogen, IB21001, US) (2014). The transference was verified by Ponceau Red (Sigma P7170, US) (2017-2019) staining for 2 min followed by washes with distilled water.

The membranes were blocked with 5% BSA diluted in TBST buffer (137 mM NaCl, 20 mM Tris, 0,1% Tween 20, pH 7.4) then incubated sequentially with anti-ATGL antibody (1:1,000 dilution in TBST with 1% BSA) overnight at 4°C. After three washes with TBST, the membranes were incubated with secondary antibodies conjugated with HRP (Kindle Biosciences, R1005 and R1006, US; diluted at 1:10,000) (2017-2019) for 1 h, followed by incubation for 1 min with the Hi/Lo Digital-ECL Western Blot Detection Kit (Kindle Biosciences, R1004, US) (2017-2019). Images of the immunoassayed blots were acquired with the Kwik Quant Imager (Kindle Biosciences, D1001, US) (2017). Then the membranes were stripped with Stripping Buffer (Thermo Fisher Scientific, 46430, US) (2017-2020) for 10 min, then washed three times with PBS and reincubated with anti-GAPDH (Genetex GT239) (2017, 2020) (1:10,000) in the same manner as above. Images were acquired as described above.

### RNA extraction and PCR

The RNA extraction was performed from both, tissues and isolated cells in culture, with the RNAeasy minikit (Qiagen, 74106, US) (2017, 2018) according to the instructions of the manufacturer, including the optional step with RNAse-free DNAse (Qiagen, 79254) (2017). RNA extracted from whole retinas was stored at −80°C until processed in the same manner as the RNA samples belonging to cultures. For RT-PCR, the samples were run in the ViiA 7 Real-Time PCR System with the QuantiTect SYBR Green (Qiagen, 204143) (2017). The relative mRNA expression was calculated using the comparative threshold method (Ct-method) with 18S for normalization. For the housekeeping gene, 18S, we used the sample at a 1:1000 dilution in RNAse-free water (Qiagen, 129112) (2017), *Pnpla*2 samples were undiluted. Each of the two biological samples were assayed for PCR in triplicate. RT-PCR primers were specifically designed to amplify the following cDNAs: *Pnpla2* Forward: (5’-TGTGGCCTCATTCCTCCTAC-3’), *Pnpla2* Reverse (5’-TGAGAATGGGGACACTGTGA-3’), *18s* Forward (5’-GGTTGATCCTGCCAGTAGC-3’),*18s* Reverse (5’-GCGACCAAAGGAACCATAAC-3’). The efficiency was approximately 90% to each pair of primers.

### Statistical analysis

Statistical analysis was performed by GraphPad Prism, using One-ANOVA, followed by the Tukey’s multiple comparison test or Dunnet’s multiple comparison test, as suggested by the software. Data are shown as mean ± S.E.M. of at least three independent experiments. Differences were considered significant at p <0.05 (*); p <0.01 (**), p< 0.001 (***), and p<0.0001 (****). When needed, the Shapiro-Wilk test was used to verify the normality of the data; as well as the ROUT (Q=1%) for outlier removal. For all experiments, the experimenters were unaware of the experimental conditions. No sample calculation was performed. No exclusion criteria were pre-determined.

## Results

### PEDF-R is present in primary retinal neurons in culture

We cultured primary neurons obtained from retinas of one day old (PN1) rats and examined the presence of the receptor for PEDF. As shown in Figure 1A, we assessed the expression of the *Pnpla2* gene in primary retinal neuronal cultures at 1, 3 and 5 days *in vitro* (DIV), and compared the *Pnpla2* expression levels with those in the whole retina, at 2, 4 and 6 post-natal (PN) days. Given that the cell cultures were from rats at PN1, the *in vitro* 1, 3 and 5 days (DIV1, DIV3 and DIV5) corresponded to their equivalent *in vivo* PN2, PN4, and PN6, respectively. RT-PCR revealed that the maximum *Pnpla2* transcription values (*Pnpla2/18S* ratio) were obtained at PN2 *in vivo* and in its equivalent, first day *in vitro* (DIV1) (Fig. 1A). As expected, the *Pnpla2* expression levels were lower for the *in vitro* than the *in vivo* cells. The *Pnpla2* transcription values in the *in vivo* retina then rapidly and significantly decreased from about 15.7 at day PN2 to nearly one third and one seventh (5.7 and 2.2) at PN4 and PN6, respectively. The ratios obtained from *in vitro* experiments, also decreased from about 4 in the first day *in vitro* (DIV1) down to nearly one third (1.25) in the fifth day *in vitro* (DIV5). Then we assessed the production and distribution of PEDF receptor (PEDF-R) protein in pure neuronal cultures. Western blot assays of neuronal cell cultures at the same intervals showed that the protein followed a similar pattern to that of the mRNA (Fig. 1B), decreasing with time in culture. Immunocytochemical evaluation of PEDF-R in retinal neurons *in vitro* revealed that it had a patched distribution in the cytoplasm around the nucleus and along the axons, colocalizing with the plasma membrane marker, wheat germ agglutinin (WGA), in both of the cell types that are known to be enriched in the cultures, amacrine and PHR neurons (Figs. 1C). A similar co-localization was observed when comparing the staining patterns of PEDF-R and another plasma membrane marker, anti Na+-K+ pump (Fig. S1). Orthogonal projection by confocal microscopy revealed that these proteins were localized mainly in the plasma membranes (Fig. 1D). We conclude that PEDF-R is present in both amacrine and photoreceptor cells, located primarily on the cell membrane. *PEDF, and the 44-mer and 17-mer fragments protected retinal neurons from cell death*.

**Figure 1.**
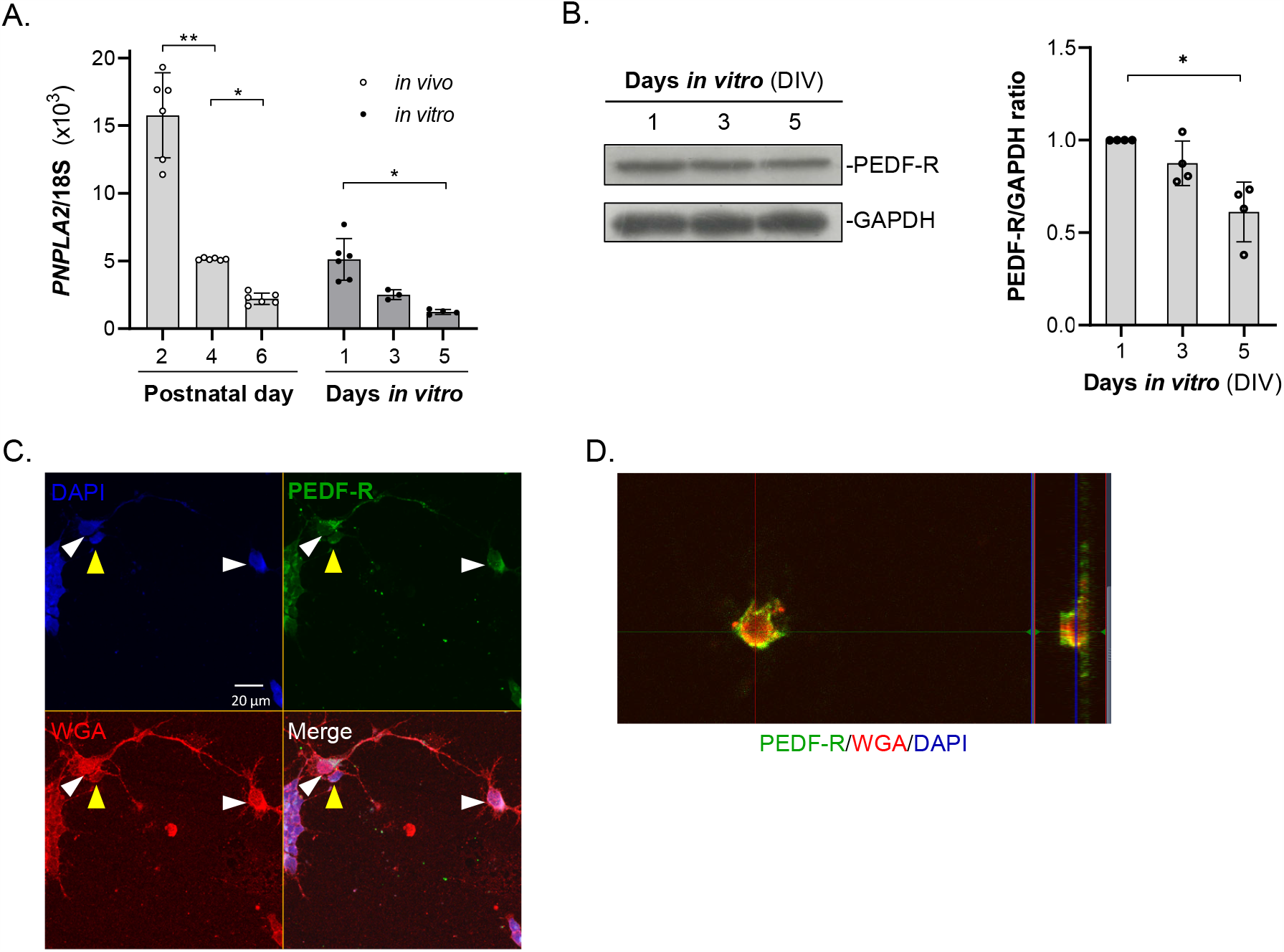
Expression of *Pnpla2* and distribution of PEDF-R. RNA expression of *Pnpla2* in retina cells in culture *in vitro* and *in vivo*. Data were collected by RT-PCR with RNA obtained from retinal cultures at DIV1, 3 and 5 and their equivalent of in vivo retinas of 2, 4- and 6-day old pups, and plotted to show the *Pnpla2*/*18s* ratios as a function of the days in vitro or age in vivo (+/- SD) (n=3; n= number of animals or number of independent cell culture preparations). Statistical analysis was performed with a One-way ANOVA with a post-hoc Tukey’s Test. * p<0.05, ** p<0.01. B) PEDF-R protein in the cell cultures determined by Western blot of protein extracts from cell cultures at DIV1 (lane 1) DIV3 (lane 2) and DIV5 (lane 3), stained with anti-PNPLA2 antibody (50 kDa band) and anti-GAPDH (38 kDa band). (n=3; n= number of independent cell culture preparations). Statistical analysis was performed with a One-way ANOVA with a post-hoc Tukey’s Test. * p<0.05. C) Photomicrograph of DIV5 retinal neuronal cultures immunostained with PEDF-R (green), the membrane marker Wheat Germ Agglutinin (red) and counterstained with DAPI (blue). Pictures show amacrine cells (white arrowheads) and PHRs (yellow arrowheads). D) Orthogonal projection of a Z-Stack showing the colocalization area of PEDF-R (Red) with the membrane marker WGA (Green), taken from the previous culture.

Previous work has shown that in the absence of trophic factors, PHRs degenerate during time in culture (*Rotstein et al., 1996; 1998*). To evaluate the potential pro-survival effects of PEDF and its derived peptides, neuronal cultures were incubated with either PEDF, the 17-mer and 44-mer (neurotrophic) peptides, or the 34-mer (antiangiogenic) peptide at DIV 2. Then cell death was determined at DIV5 by measuring cells that had nuclei labeled with propidium iodide (PI) and terminal deoxyribonucleotidyl transferase (TDT)-mediated dUTP-digoxigenin nick end labeling (TUNEL). PI, a small fluorescent molecule that binds DNA but cannot passively traverse into cells with an intact plasma membrane, was used to discriminate dead cells, in which plasma membranes become permeable regardless of the mechanism of death, from viable cells with intact membranes. While control cultures (treated with vehicle alone) had about 29% of PI-positive cells, those treated with 10 nM PEDF, 44-mer peptide or 17-mer peptide had lower percentage of non-viable cells (13%, 15%, and 16%, respectively) (Fig. 2B). In contrast, when the cultures were treated with effectors preincubated with 100 nM of blocking P1 peptide, the percentage reverted (29%, 30%, and 27%, respectively) (Fig. 2B).

**Figure 2.**
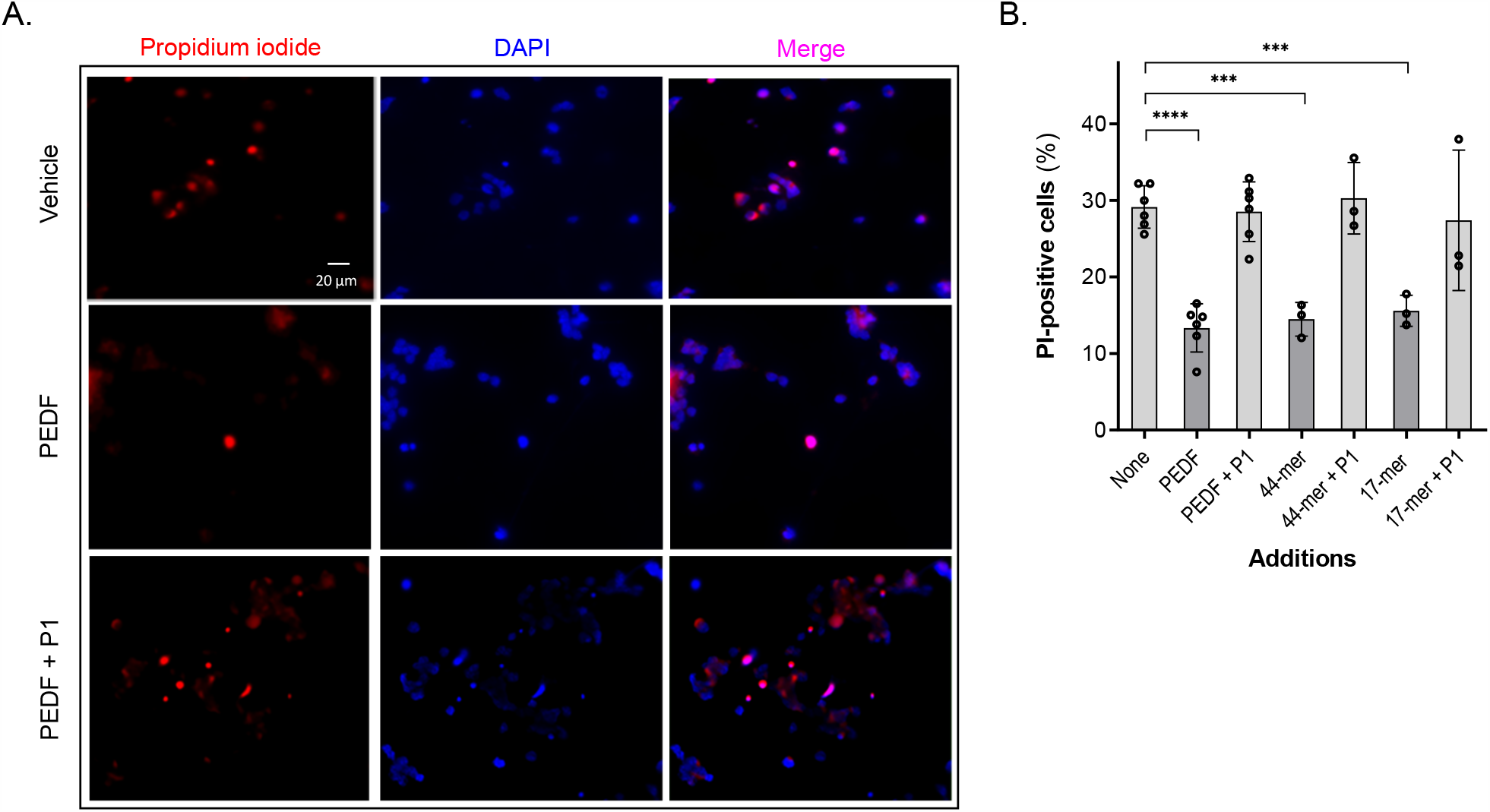
Effects of PEDF and its derived peptides in the prevention of cell death. A) Fluorescence photomicrographs of 5-day cultures supplemented with vehicle (control)-, PEDF-, and PEDF + P1, showing PI (red) stained neurons (arrows), and counterstained with DAPI (blue) (Bar: 10 µm). B) Percentages of PI-positive nuclei in cultures treated without (None) or with PEDF; and PEDF-derived peptides, preincubated or not with the blocking peptide, P1, as indicated in the x-axis. Statistical analysis was performed using a One-way ANOVA with a post-hoc Dunnett’s test. *** p<0.001, **** p<0.0001 (n=6; n= number of independent cell culture preparations).

TUNEL of free 3’-hydroxyl termini of double-stranded DNA fragments *in situ* was performed to detect DNA damage in cells. Control cultures had approximately 8-12% of TUNEL-positive cells, and most of them displayed fragmented or pyknotic nuclei (Fig. 3). Additions of PEDF, 44-mer or 17-mer peptides (10 nM) decreased the number of TUNEL-positive cells, being 62 ± 6.2%, 45 ± 6.5%, 59 ± 0.9%, respectively, of that of the control value (Fig. 3B). In all cases, preincubation of the effectors with a 10-fold molar excess of the blocking P1 peptide abolished these effects. The antiangiogenic 34-mer peptide and the P1 peptide alone had no protective effect on these cultures (Fig. 3B).

**Figure 3.**
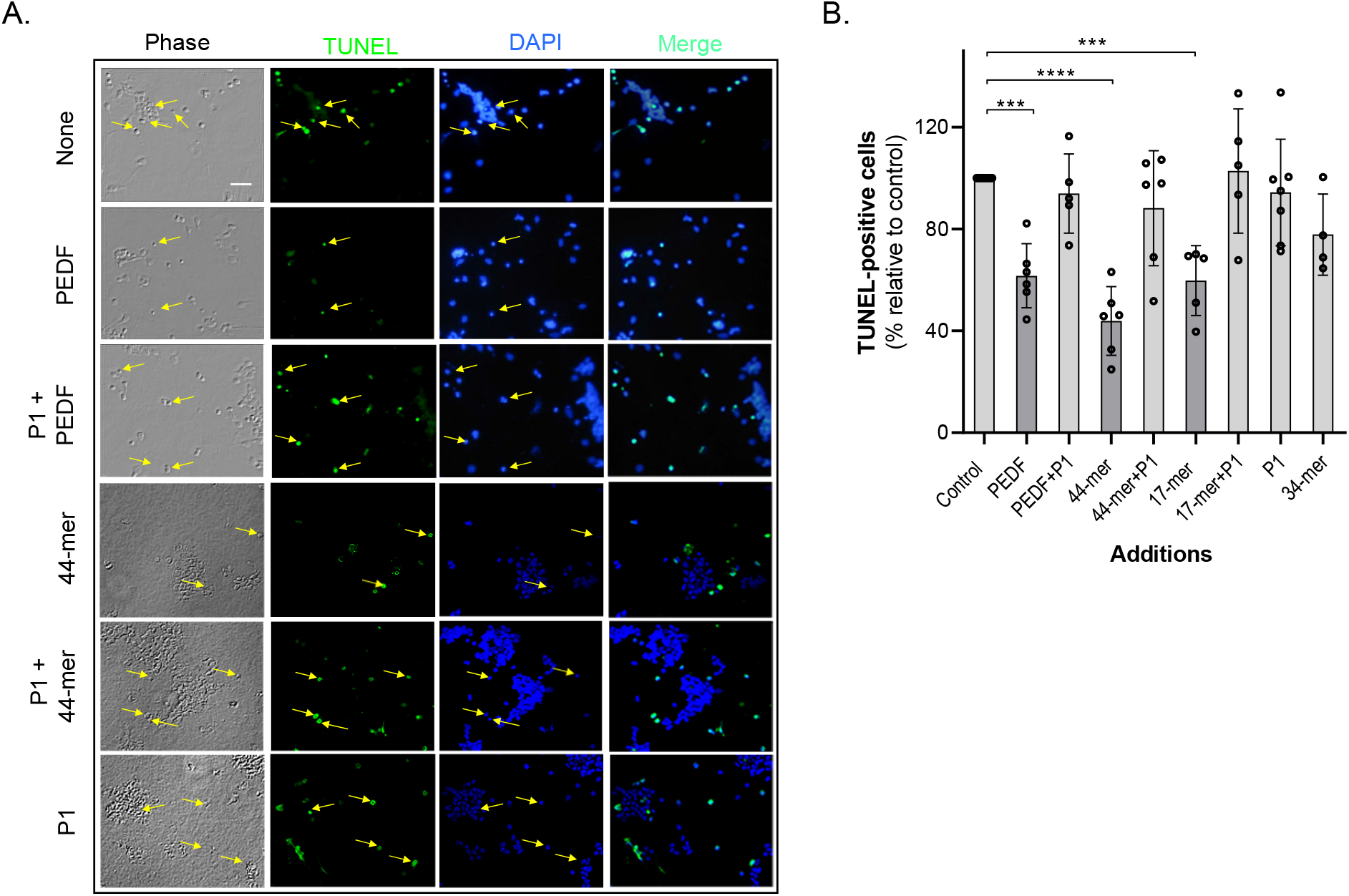
Effects of PEDF and its derived peptides in apoptosis prevention. A) Photomicrographs of DIV5 cultures showing TUNEL (green label) positive neurons (arrows), and counterstained with DAPI (blue) (Bar: 20 µm); B) Percentages of TUNEL-positive nuclei relative to control samples. Statistical analysis was performed using a One-way ANOVA with a post-hoc Dunnett’s test. ** p<0.01, *** p<0.001, **** p<0.0001 (n=5; n= number of independent cell culture preparations).

Flow Cytometry analysis to evaluate neuronal death in cultures treated with PEDF and the peptides was also performed. Fig. 4 shows that PEDF decreased the percentage of TUNEL-positive cells (Q2+Q3) by half, from 39.3% in controls to 19.2%. This effect was reversed when the cultures were pre-incubated with atglistatin, an inhibitor of the enzymatic activity of PEDF-R prior to PEDF addition. In contrast, control conditions and 34-mer treated-cultures had similar percentages of TUNEL-positive cells (39.3% and 37.0%, respectively), as indicated by BrdU intensity.

**Figure 4.**
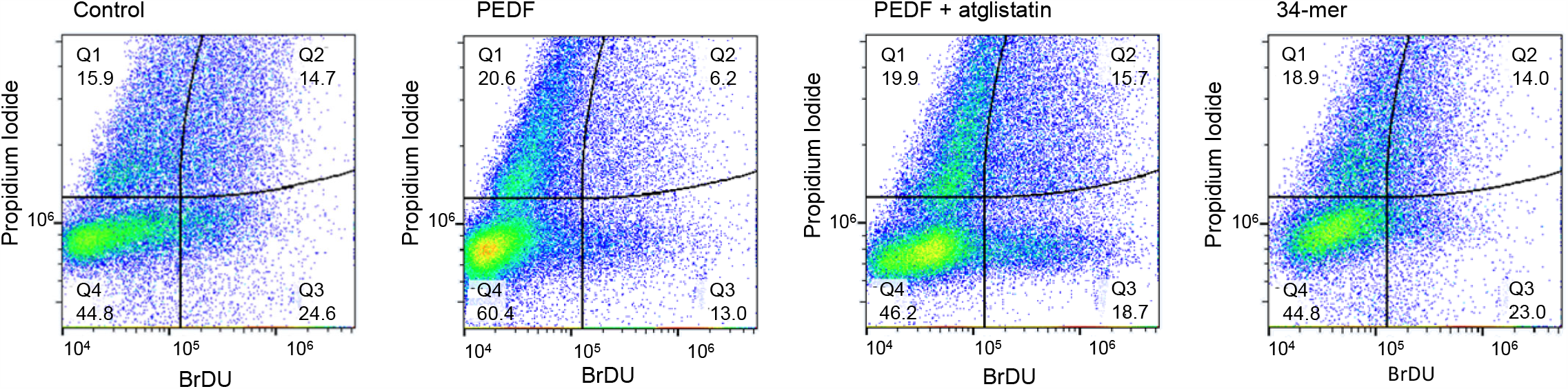
Flow Cytometry analysis of cell death by TUNEL of DIV5 cultures treated without (control), or with PEDF; PEDF + Atglistatin; or 34.-mer peptide and determined as BrDU intensity (X axis), and propidium iodide (Y axis).

Altogether, these results show that PEDF and its neurotrophic fragments, 44-mer and 17-mer, specifically prevented neuronal retinal cell death and DNA damage, implicating the PEDF/PEDF-R axis in promoting retinal cell survival.

### Flow cytometry reveals two distinct populations of cells

Flow Cytometry analysis of the cultures revealed the presence of two cell populations (Figs. 5A-5B). One of them, the low forward scatter (low FSC), had cells of smaller size representing 40.9% of the total cells, and the second one, the high FSC, had larger cells representing 55.5% of the total cells for the control condition (Fig. 5A).The amount of events for these groups in the PEDF-treated condition also had two population, amounting to 25.3% and 52.5%, for low and high FSC, respectively (Fig. 5B) The low FSC and high FSC cell populations likely corresponded to PHRs and amacrine neurons, respectively, which differ in cell size and are the main neuronal cell types previously described in the primary retinal cell cultures used here (Politi et al. 1988).

**Figure 5.**
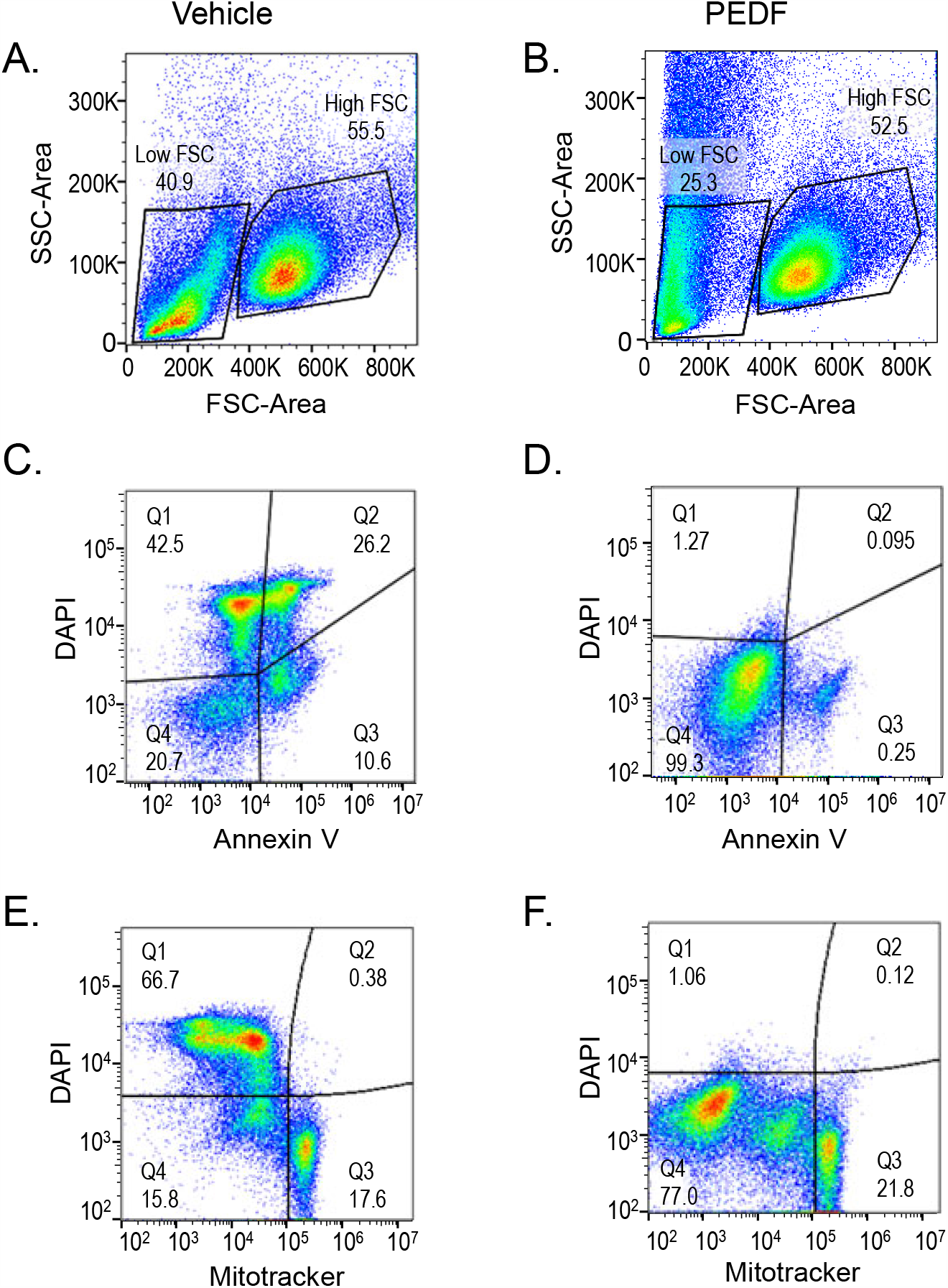
Flow Cytometry assays of two neuronal populations at 5 and 7 days in vitro. A, B): Forward-Side Scattering plots of DIV5 Vehicle-(Control), and PEDF-treated cultures, respectively, showing the low- and high-forward scatter (FSC) neuronal populations (presumptive PHRs and amacrine cells, respectively): C, D): DAPI (Y axis) and Annexin V (X axis) labeled cells, and (E, F) DAPI (Y axis) and Mitotracker (X axis) plots of the low-FSC population.

### PEDF inhibitory effects on early events of cell death in the low FSC cell population

Annexin V, a phospholipid-binding protein, was used to detect redistribution of phosphatidyl serine that occurs from the inner to the outer leaflet of plasma membrane lipid bilayers during early events of cell death. DAPI, a fluorescent stain that binds strongly to A-T-rich regions in DNA, was also used as it inefficiently traverses intact cell membranes and therefore, preferentially stains dead cells. Analysis of the low FSC population, the PHRs, under control conditions indicated that about 21% of cells were negative for Annexin V and DAPI staining (Fig. 5C, Q4) while the remaining 79% (Fig. 5C, Q1-Q3) were positive, implying that this population underwent cell death. In contrast, when the cultures were treated with PEDF nearly 92% of cells were negative for Annexin V and DAPI staining (Fig. 5D, Q4), indicating PEDF-mediated protection against cell death. These results imply that PEDF prevented phosphatidyl serine flipping and loss of structural integrity of the plasma membrane consistent with inhibition of early events of PHR cell death.

### PEDF and the 44-mer fragment selectively increased mitochondrial membrane potential in PHRs

To evaluate mitochondrial involvement in PEDF-induced cell survival, we determined Mitotracker-positive cells among the cells in the low FSC population of presumptive PHRs by flow cytometry. MitoTracker Red CMXRos, a cell permeable probe that contains a mildly thiol-reactive chloromethyl moiety, was used to stain mitochondria in live cells because it passively diffuses across the plasma membrane to accumulate in active mitochondria in a membrane potential-dependent fashion. The results showed that, on the one hand, 17.6% of cells in controls exhibited intense Mitotracker staining and were negative for DAPI staining (Fig. 5E, Q3). This observation indicated that a few cells preserved healthy mitochondria and intact plasma membrane. On the other hand, 66.7% of cells showed a weak Mitotracker staining and strong DAPI staining (Fig. 5E, Q1), indicating that these cells had their mitochondrial function and cell membrane compromised. In contrast, when the cultures were treated with PEDF about 22% of presumptive PHRs were heavily stained with Mitotracker (Fig. 5F, Q3) and none of these cells were DAPI positive (Fig. 5F, Q1, Q2), indicating that PEDF preserved mitochondrial function.

### Effects of PEDF on the population of the high FSC cell population

The population of larger sized cells, presumptively amacrine neurons, were negatively stained for DAPI and Annexin V under both control conditions and PEDF-treated conditions (Figs. 6A, 6B, Q4), indicating that they had intact cell membranes. Nearly all these neurons exhibited intense Mitotracker staining, indicating that they had healthy mitochondria even in the absence or presence of PEDF (Figs. 6C and 6D, Q3). In this regard, it is important to mention that the chemically defined medium used for these cultures contains insulin, which we have previously demonstrated is a trophic factor for these neurons (Politi et al., 2001b).

**Figure 6.**
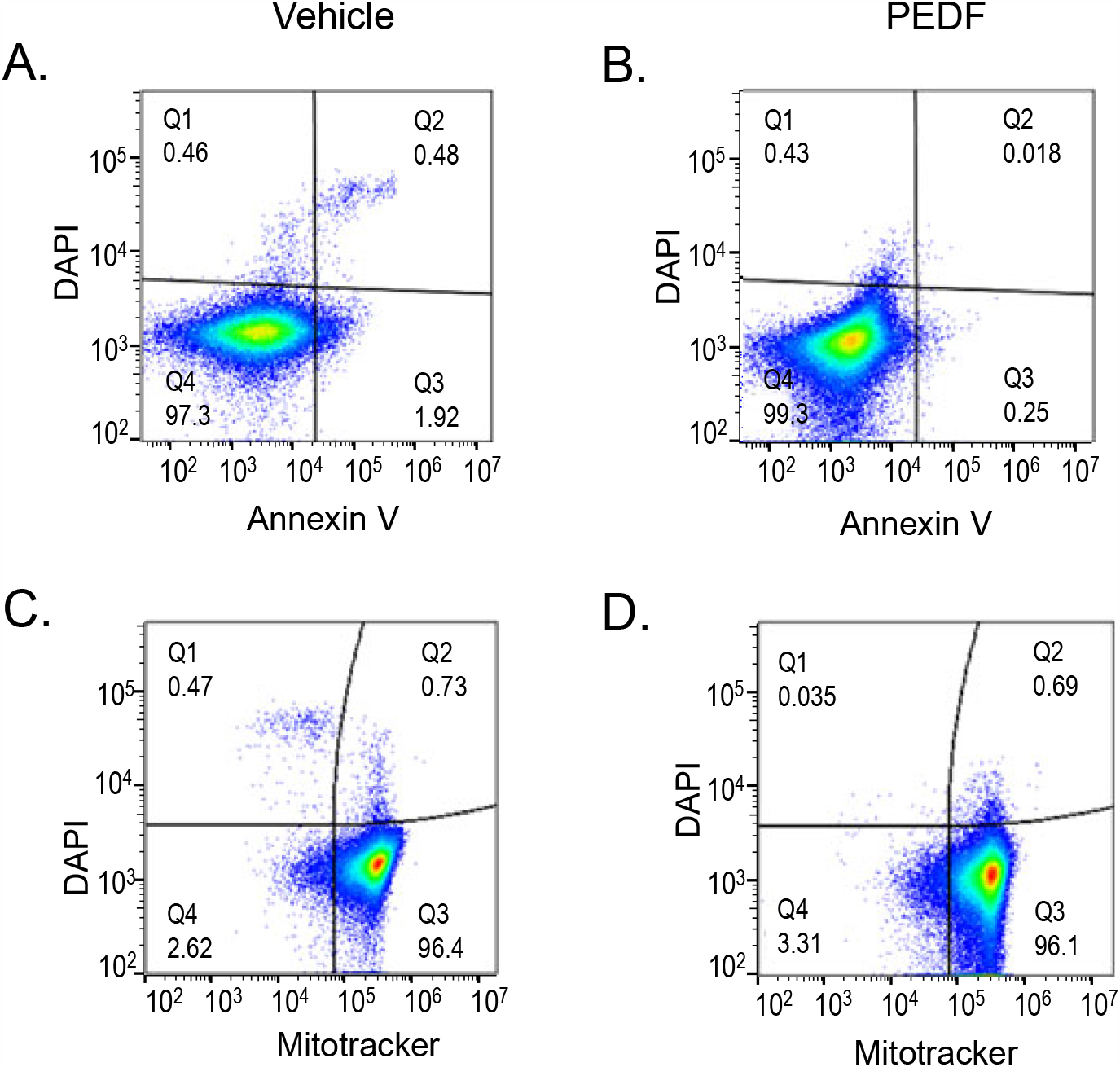
Flow Cytometry assays of high FSC cell populations at DIV5. Plots of the high-FSC population showing DAPI and Annexin V (A and B) and DAPI and Mitotracker (C and D) labeled cells.

### Survival effects of PEDF on cultured PHR cells at DIV7

At DIV7, the FSC/SSC analysis by Flow Cytometry, revealed that in controls, the low FSC Gate population of PHRs, almost completely disappeared, representing less than 1% (Fig. 8A). However, incubation of the cultures with either PEDF or the 44-mer peptide increased these values to 9.6% and 13.5% respectively (Figs. 7B, 7D), indicating that they partially preserved the population of presumptive PHRs. Preincubation of the cultures with atglistatin gave similar values to control cultures (compare Figs. 7B and 7C). Thus, the requirement of PEDF for PHR survival was evident at day DIV7.

**Figure 7.**
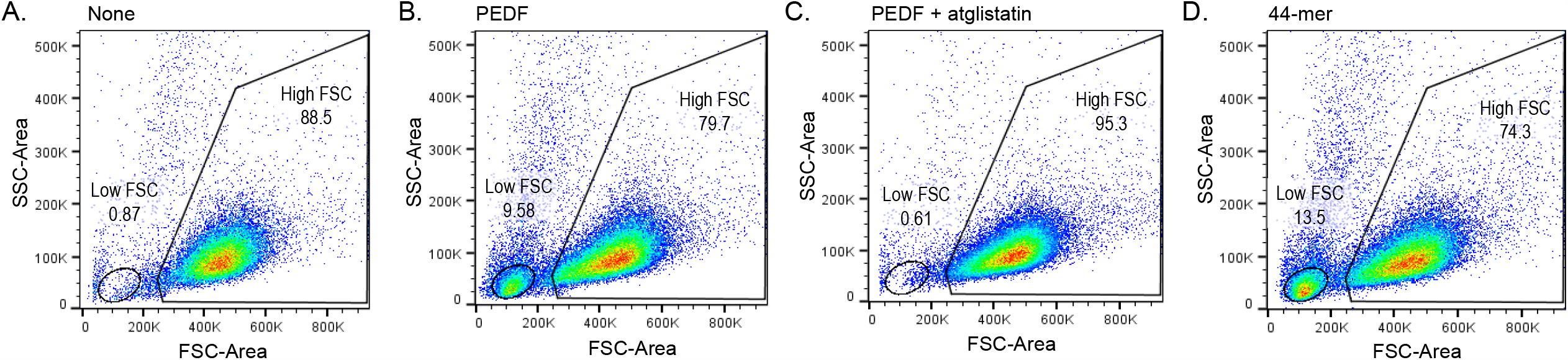
Flow Cytometry assays of two neuronal populations at 5 and 7 days in vitro. Forward-Side Scattering plots of DIV7 cultures showing the low-, and high-FSC neuronal populations in control (A); PEDF (B); PEDF plus Atglistatin (C); and 44-mer (D); and treated cultures.

**Figure 8.**
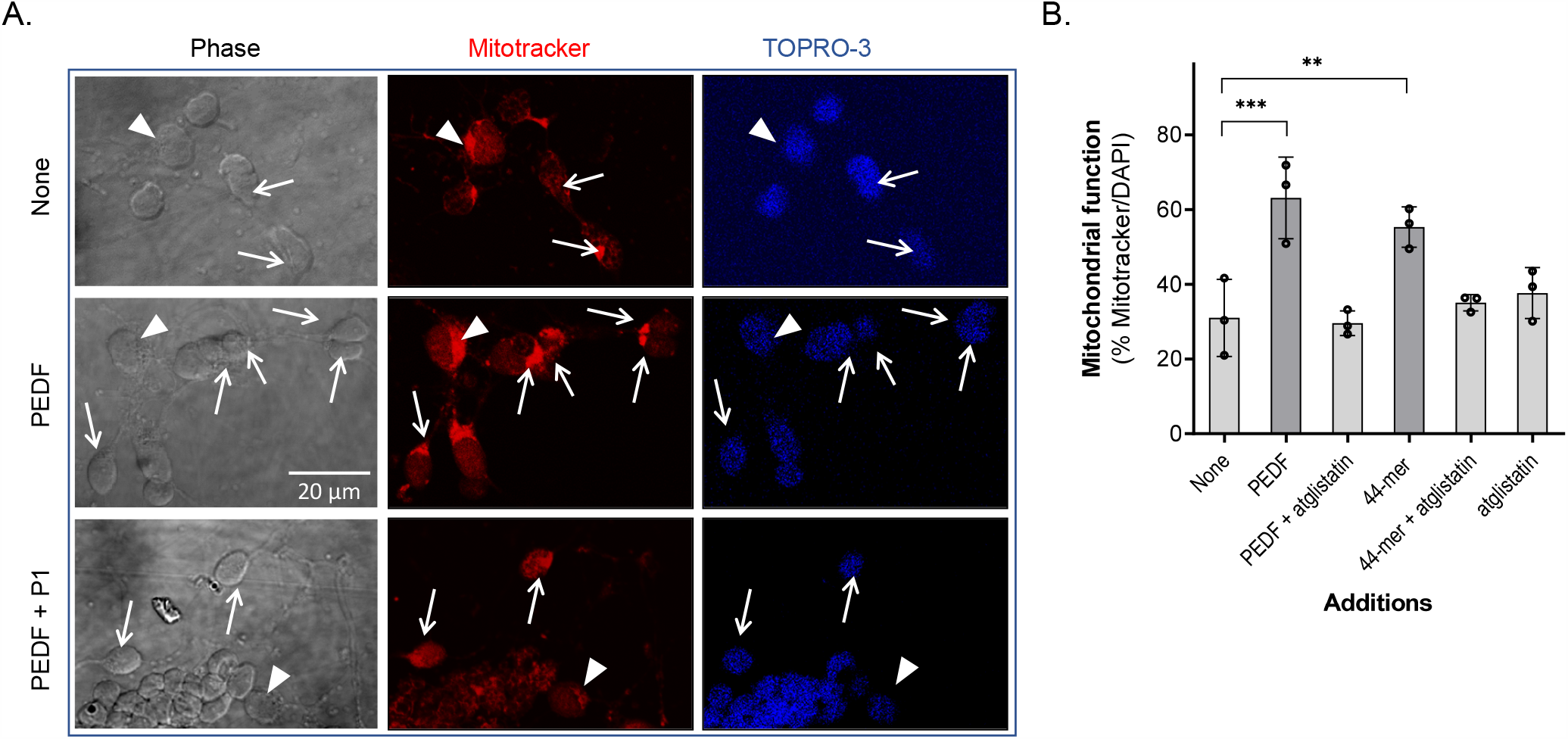
Effects of PEDF and its derived peptides in the preservation of mitochondrial activity. A) Phase and fluorescence photomicrographs of DIV5 vehicle-(control), PEDF-, and PEDF+P1-treated cultures showing Mitotracker (red) and TOPRO-3 (blue) stained cells. Arrowhead points to amacrine and arrows mark PHR neurons with active mitochondria. Note that amacrine neurons showed brilliant (active) mitochondria regardless PEDF supplementation, while PHRs depend on PEDF to retain their active mitochondria. Both neuronal cell types were identified as indicated in Materials and Methods. (Bar: 20 µM). B) Percentage of PHRs bearing active mitochondria in cultures treated with HBSS (control); PEDF; with and without P1. Statistical analysis was performed using a One-way ANOVA with a post-hoc Dunnett’s test. ** p<0.01, *** p<0.001 (n=3 n= number of independent cell culture preparations).

### Cytochemical examination of mitochondria of neuronal retinal cells treated with PEDF

Neuronal cultures stained with MitoTracker showed brilliant punctate red dots in the cytoplasmic areas around the nucleus and in the axon hillock (Fig. 8A), illustrating preservation of mitochondrial membrane potential in both, amacrine and PHR neurons (arrows in Fig. 8A). Amacrine neurons and PHR neurons were identified as indicated in Materials and Methods. In control cultures, 31% of PHRs (White arrows) retained active mitochondria by day DIV5, and this value doubled (63%) when cultures were incubated with PEDF or with 44-mer (Fig 8B). The increases in Mitotracker-positive cells were blocked by the inhibitor atglistatin and by pre-incubation with blocking P1 peptide (Fig. 8A). The amacrine neurons retained active mitochondria, regardless of the absence or presence of PEDF or its derived factors (cells labeled with white arrowheads in Fig 8A).

### PEDF promoted opsin apical localization on PHRs

At DIV5 and in the absence of neurotrophic factor, most rhodopsin-positive PHRs showed a diffuse opsin distribution along the cell membranes, both in the cell body and axons, reflecting an immature stage of differentiation (Figs. 9A and S2A). During PHR development and differentiation, rhodopsin tends to gradually fade away from the axons (Fig. S2B), until it is solely located in the cell body. Later, rhodopsin starts to polarize toward the apical region (Fig. S2C), where it finally constitutes a rhodopsin-positive structure (Fig. S2D). However, under control conditions at DIV5, only 28% of PHRs reached this rather mature state of differentiation (Fig. 9B). Remarkably, addition of PEDF dramatically increased the percentage of PHRs having rhodopsin positive apical processes to about 72.5% (Fig. 9B). The effects of the 17-mer and 44-mer fragments mimicked those of PEDF and were blocked by atglistatin (Fig. 9B). These results imply that PEDF and the 44-mer and 17-mer fragments promoted PHR differentiation and proper rhodopsin localization. However, the number of PHRs containing rhodopsin remained the same in control cultures as in those with PEDF and peptides (Fig. 9C).

**Figure 9.**
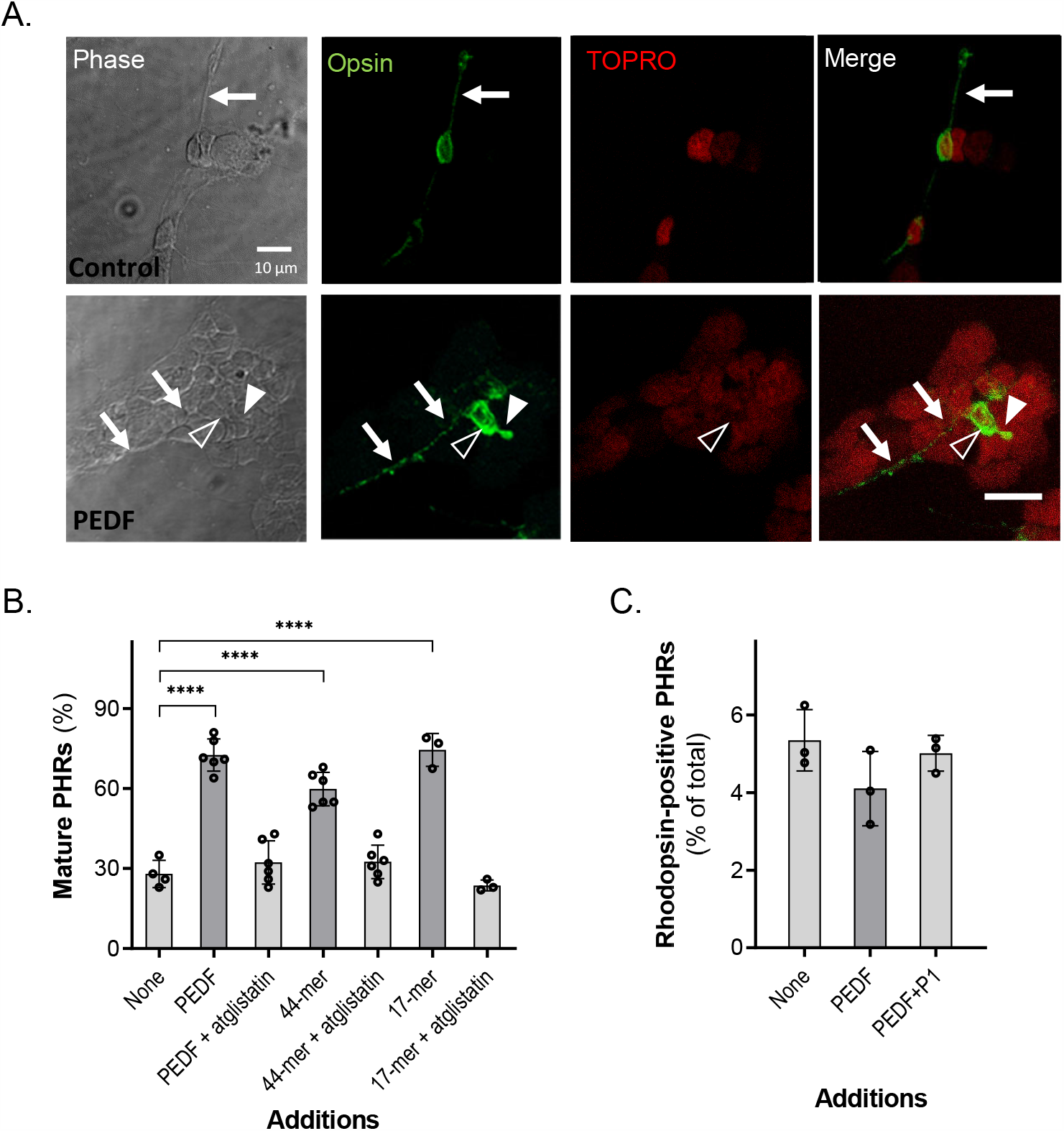
Effects of PEDF and its derived peptides on rhodopsin distribution in PHRs during development. A) Differential interference contrast (DIC) and fluorescence photomicrographs of DIV5 cultures showing opsin (green) distribution, counterstained with TOPRO-3 (red) in control (no PEDF) and PEDF-treated PHRs. Note that in control cultures, rhodopsin is homogenously distributed in both the cell body and neurites (arrows), while in PEDF-treated conditions, rhodopsin is concentrated in the cell body and in the outer segment-like structure (arrowheads) (Bar: 10 µm); B) Percentages of rhodopsin positive PHRs labelled only in the cell body. Statistical analysis was performed using a One-way ANOVA with a post-hoc Dunnett’s test. **** p<0.0001 (n=8). C) Percentages of rhodopsin positive PHRs in control; PEDF and PEDF + P1 treated cultures (n=6; n= number of independent cell culture preparations).

### The 44-mer and 17-mer fragments promoted neurite outgrowth

Like PEDF, the 44-mer and 17-mer peptides also exhibited an effect on neurite outgrowth, which was driven particularly by amacrine neurons, as shown by immunocytochemistry using antibodies against βIII Tubulin and HPC-1, biomarkers for neurons and early amacrine cell development, respectively (Fig. 10A). PEDF, and the 44-mer and 17-mer fragments increased the average of amacrine neurons bearing axons with lengths of 4-fold their cell diameters to 3.3, 4.0 and 3.6 length/diameter ratio, respectively, from 2.8 in length/diameter ratio for control cultures treated with PBS as vehicle (Fig. 10B). The inhibitor of PEDF-R atglistatin decreased the average of neurite length/diameter ratios of neurons treated with PEDF, 44-mer and 17-mer to 2.8, 2.9 and 3.1 in length/diameter ratio, respectively. Addition of DMSO, the vehicle used for atglistatin, did not alter the PEDF efficacy, and the 34-mer did not induce neurite outgrowth. Moreover, cultures incubated with 10-fold molar excess of the blocking P1 peptide over PEDF showed no effect on neurite outgrowth, indicating that the longest neurites corresponded to those of PEDF-treated cultures (Fig. S3). These observations demonstrated that the 44-mer and 17-mer fragments, like PEDF, specifically promoted neurite outgrowth in amacrine neurons in culture.

**Figure 10.**
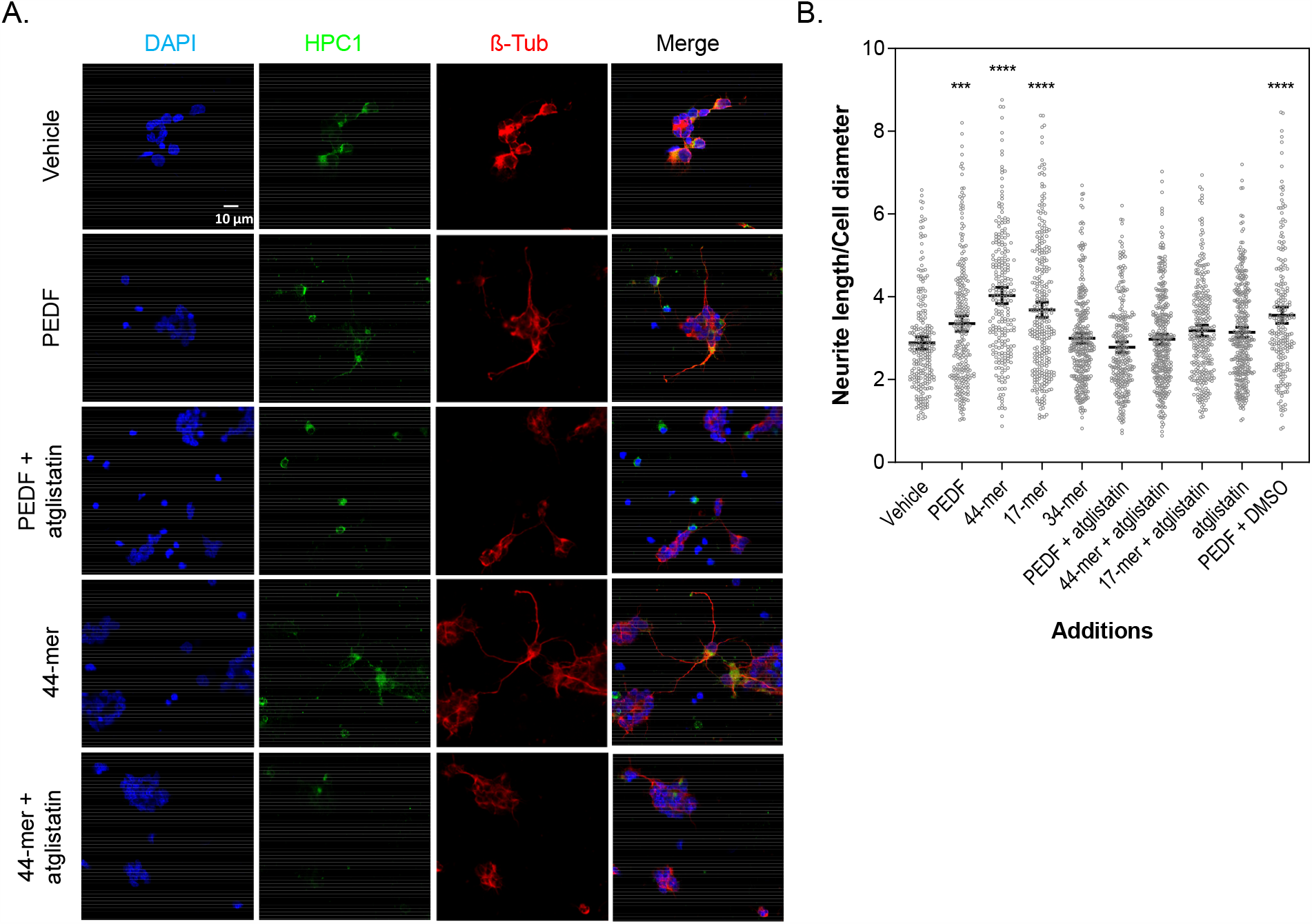
Effects of PEDF and its derived peptides on neurite outgrowth. A) Fluorescence photomicrographs of DIV5 cultures showing HPC-1 positive (green) amacrine neurons, labeled with ß-Tubulin (Red) and counterstained with DAPI (blue), for neurite outgrowth in control; PEDF, 44-mer; and PEDF plus atglistatin-treated cultures. (Bar: 10 µm). B) Dot-plot showing neurite length measurements under different treatments, as indiacted in the x-axis. Statistical analysis was performed using a One-way ANOVA with a post-hoc Tukey’s test. *** p<0.001, **** p<0.0001, respect to control (None). (n=3; n= number of independent cell culture preparations)

## Discussion

In this study, we examined the neurotrophic effects of PEDF fragments in a model of cultured primary retinal neurons that that undergo programmed cell death in the absence of trophic factors *in vitro*. The results show that full-length human PEDF protein and the small 44-mer and 17-mer peptides (78-121 and 98-114 residue positions of human PEDF, respectively) prevent cell death and promote differentiation of rat retinal PHRs and amacrine neurons *in vitro*. Moreover, the PEDF-mediated effects on PHR cell survival and differentiation are associated to prevention of plasma membrane disruption, phosphatidyl serine flipping, DNA damage and preservation of mitochondrial membrane potential, and implying that PEDF inhibits early in the cell death proces.. In addition to promoting apical localization of opsin in PHRs the 17-mer and 44-mer peptides selectively stimulate axonal outgrowth in retinal amacrine neurons. Noteworthy, the antiangiogenic peptide 34-mer (44-77 positions) lacked these activities. Interestingly, the distribution of PEDF receptors (PEDF-R) in plasma membranes explain the diverse activities mediated by PEDF/PEDF-R interactions in both PHRs and amacrine neurons, which agrees with a cell-context dependency of the pleiotropic functions of PEDF mediated by PEDF-R activation.

To the best of our knowledge, this is the first report demonstrating the neurotrophic activities of PEDF in a primary cell culture system for the retina. Previous studies have demonstrated PEDF neurotrophic activities on retinal ganglion cells in culture using mixed cell cultures enriched in this cell type from adult rat retina or retinas from 7 day-old mice. PEDF protects retinal ganglion cells from glutamate- and trophic factor withdrawal-mediated cytotoxicity (*Pang et al. 2007*) and hypoxic insult (*Unterlauft et al. 2012*) and promotes axon growth in these cells (*Vigneswara et al 2013*). Regarding the cultures in the present study, most PHRs undergo programmed cell death at the time of synaptogenesis and during development of retinal neurons in culture, which is equivalent to the biological process of their counterparts in the native retina *in vivo*. To prevent this programmed cell death, PHRs become strongly dependent on the supply of trophic factors and cells that fail in obtaining them eventually die (*Adler 1986; Watanabe et al. 1992*). PEDF is one of these factors, and plays a critical role as a survival-promoting factor for PHRs *in vivo* (*Cayouette et al 1999; Murakami et al. 2008; Kenealey et al 2015; Comitato et al, 2018; Hernández-Pinto et al, 2019; Chen et al 2019; Miyazaki et al 2008*) and *in vitro*, as demonstrated here.

PEDF receptors are widely distributed on the cell surface of both PHRs and amacrine neurons at their early stages of development (Fig. 1). This observation suggests a likely extensive and cell selective role for PEDF during retina development. While PEDF preserves mitochondrial activity and cell survival in PHRs, it promotes axonal development in amacrine neurons, an outgrowth effect that is not observed in PHRs. However, given that amacrine neurons do not die in these cultures, it cannot be ascertained whether PEDF acts in neuroprotecting them. Noteworthy, the neurotrophic fragments of PEDF, 17-mer and 44-mer, have the same effects as PEDF. Therefore, PEDF appears to play a critical role in preventing PHR cell death and orchestrating the development and differentiation of the retina and this role can be almost exclusively ascribed to its neurotrophic fragments.

The present study provides insight into a molecular mechanism of action for PEDF on PHR development and amacrine neurite outgrowth. The specific interaction of the neurotrophic domain of PEDF with PEDF-R occurs in the primary neuronal retinal cultures. This conclusion is based on the observed neurotrophic effects of the 44-mer and 17-mer peptides in combination with the attenuation of these effects by blocking the PEDF/PEDF-R interactions (use of blocking P1 peptide) and inhibiting the PEDF-R enzymatic activity (use of atglistatin). The fact that the neurotrophic 44-mer and 17-mer peptides have PEDF-R binding affinity (*Kenealey et al 2015*) and the unrelated 34-mer peptide does not, points to the specific requirement of the 17-mer region of PEDF for binding and activity (*Kenealey et al 2015*; Figs 2-7, S1). Thus, PEDF action requires the binding to and stimulation of PEDF-R in PHRs and amacrine cells, resembling the PEDF dependency on PEDF-R activation for survival of the rat retinal R28 cell line (*Subramanian et al 2013*).

Interestingly, the effects of PEDF on PHR survival and differentiation are similar to those exerted by DHA (*Garelli et al., 2006*). These similarities are not surprising given that the PEDF-R has a phospholipase A activity, which PEDF binding enhances its hydrolysis of phospholipids to release fatty acids, among them DHA (*Subramanian et al., 2012; Pham et al., 2017*). DHA preserves mitochondrial functionality, prevents apoptosis and promotes differentiation of PHRs (*Rotstein et al., 1996; 1997; 2003; Politi et al 2001, German et al, 2006; 2013*) resembling the effects of PEDF. Furthermore, 4-bromoenol lactone, an inhibitor of calcium-independent phospholipase A2, blocks the DHA-mediated prevention of oxidative stress-induced apoptosis of PHRs (*German et al., 2013*). Interestingly, 4-bromoenol lactone also inhibits PEDF-R and blocks the PEDF-mediated cytoprotective activity and induction of antiapoptotic *Bcl-2* (B-cell lymphoma 2) gene expression in R28 cells (*Notari et al 2006 Subramanian et al 2013*). Thus, it appears that PEDF and DHA are the two faces of the same coin, given that either by adding DHA or by activating PEDF-R, they promote similar survival and differentiation effects on PHRs. These similarities suggest that PEDF activation of PEDF-R leads to DHA release from phospholipids, becoming the biological mediator of the PEDF/PEDF-R interactions, decreasing death and promoting differentiation in PHRs. This hypothesis requires further study. Lipidomics and proteomics profiles of the primary rat retinal cultures treated with PEDF peptides will be instrumental to address the identification of bioactive lipids and proteins of the PEDF actions in PHRs.

In summary, the findings demonstrate that the use of primary neuronal retinal cell cultures are useful to gain understanding of the individual effects of the multimodal PEDF on the diverse cell population of the retina, as shown here for PHRs and amacrine cells. In addition, the findings establish the neurotrophic PEDF peptides as neuronal guardians for the retina, highlighting their potential as promoters of retinal differentiation, and inhibitors of retina cell death and its blinding consequences.

## Abbreviations

DIV: Days In Vitro
DAPI: 4′,6-diamidino-2-phenylindole
DHA: Docosahexaenoic Acid
FSC: Forward-Scatter
PEDF: Pigmented epithelium-derived factor
PEDF-R: PEDF Receptor
PI: Propidium Iodide
PN: Postnatal
PHR: Photoreceptor
Pnpla2: Patatin-like phospholipase-2
SSC: Side-Scatter

## Acknowledgements

This work was supported in part by the Intramural Research Program of the National Eye Institute, NIH, U.S.A. [Project #EY000306], and by the National Research Council of Argentina (CONICET): PIP 1122015, to LP and National Agency for Science and Technology (ANPCYT): PICT 2016-0353, to LP), American Society for Biochemistry and Molecular Biology USA (PROLAB award to GM). We thank laboratory members for continuous encouragement and ideas and the staff of the Politi’s lab and NEI animal facility, Confocal Microscopy Services and Biological Imaging Core and Flow Cytometry Core facilities for technical support, Marcela Vera, for sharing their expertise with western blotting, and Dr. Robert Molday (University of British Columbia, Canada) for generously providing antibody to rhodopsin.

## Supplementary information

**Figure S1.**
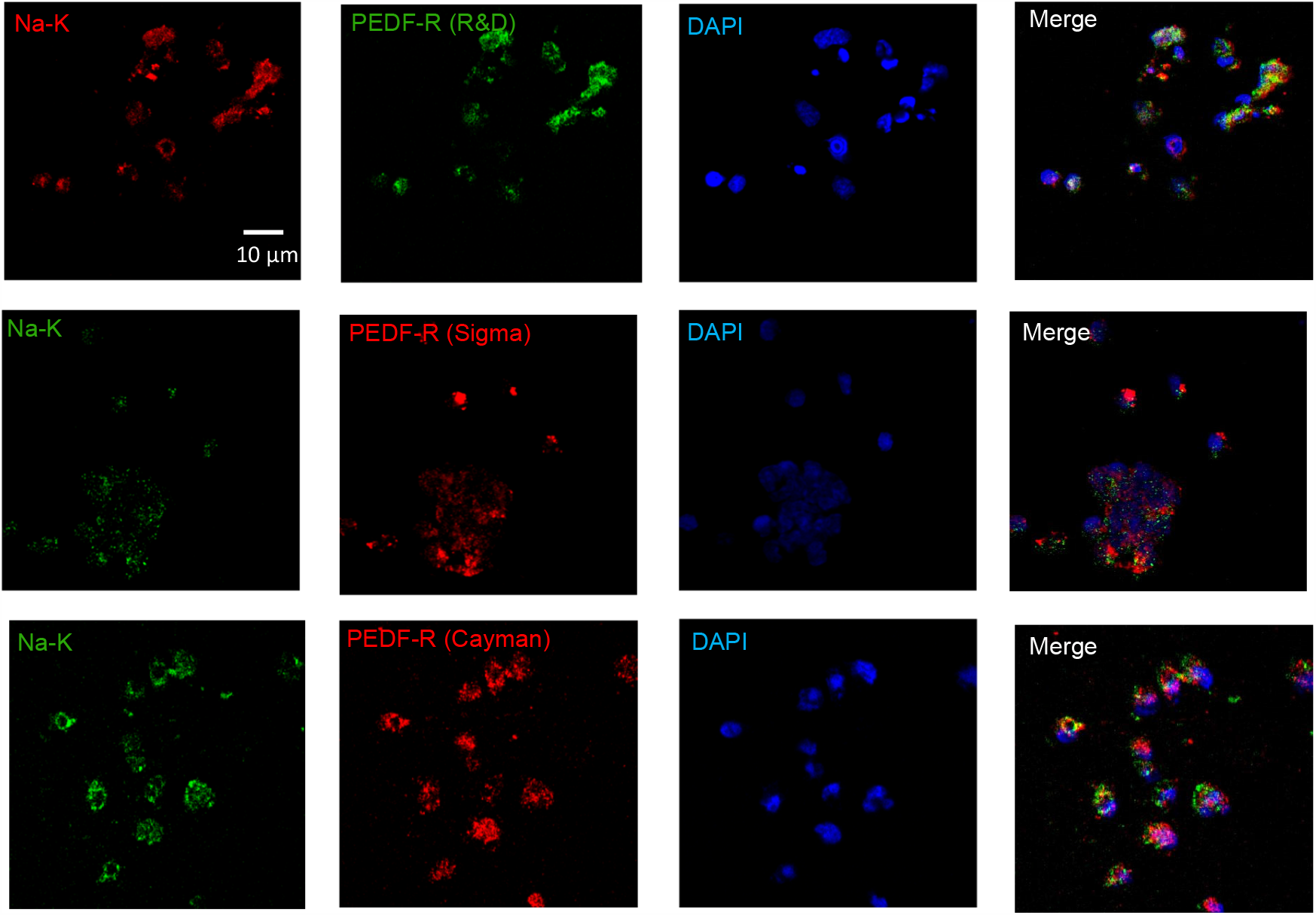
Co-expression of PEDF-R and the Na^+^-K^+^ Pump. Expression of PEDF-R and the membrane marker, Na^+^-K^+^ Pump, in 5-day neuronal cultures. PEDF-R expression was tested with antibodies from three different manufacturers (R&D, Sigma-Millipore and Cayman), which co-expressed with Na^+^-K^+^ pump (Bar 10 µm).

**Figure S2.**
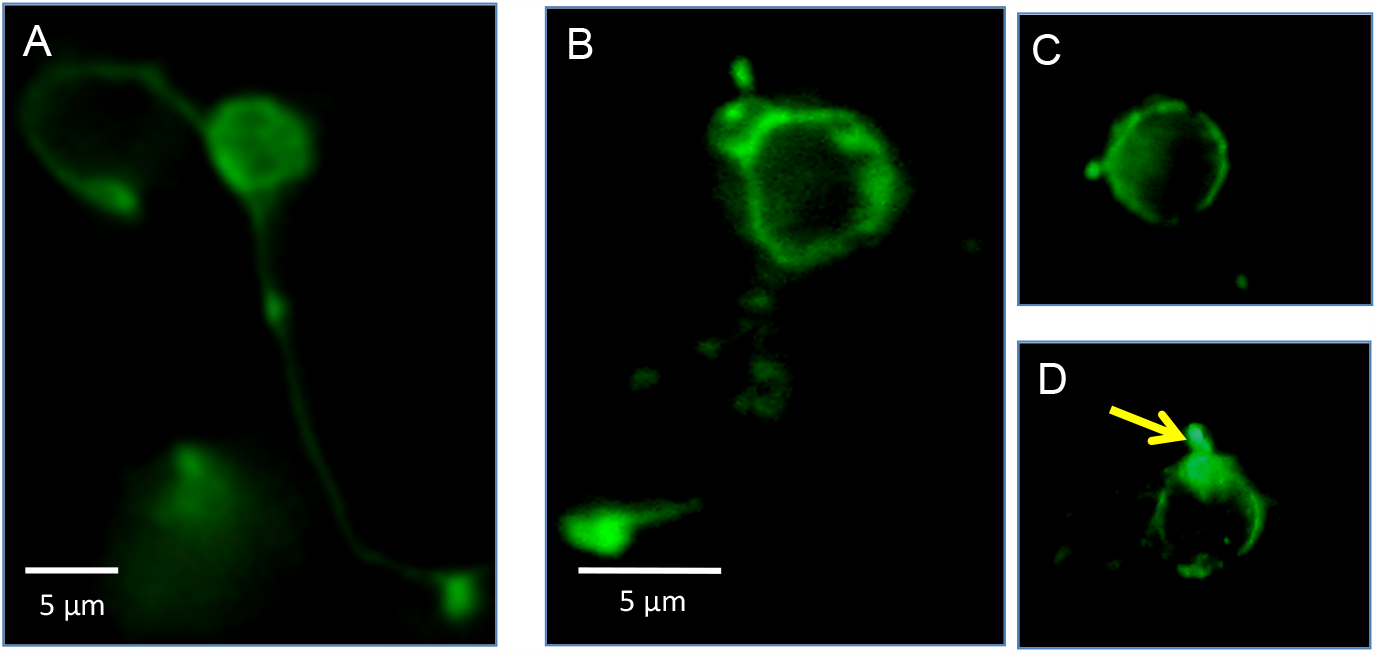
Different stages of PHR development and their respective rhodopsin distribution. Starting from a homogenous, diffuse distribution (1), which starts to fade away from the axons and concentrate on the cell body (2), from which further polarizes towards one end of the cell (3) and forms a strongly rhodopsin+ protrusion, similar to an outer segment (4) (Bar: 5 µm).

**Figure S3.**
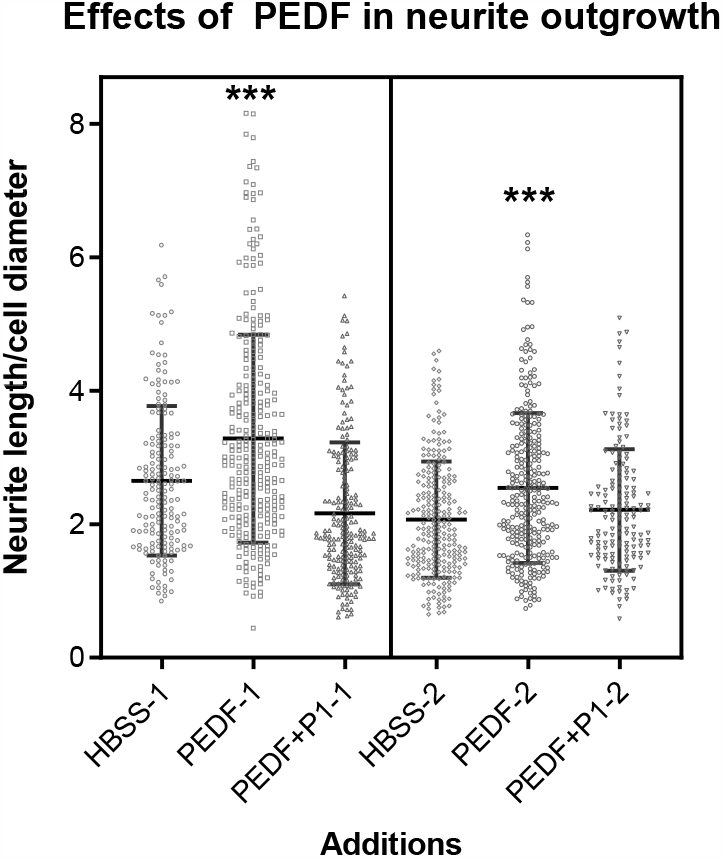
Effects of PEDF and its derived peptides on neurite outgrowth. Additional dot-plots showing neurite length measurements under different treatments. Statistical analysis was performed using a One-way ANOVA with a post-hoc Tukey’s test. *** p<0.001, respect to their respective control (HBSS-1/2) (n=3; n= number of independent cell culture preparations)

**Table S1.**
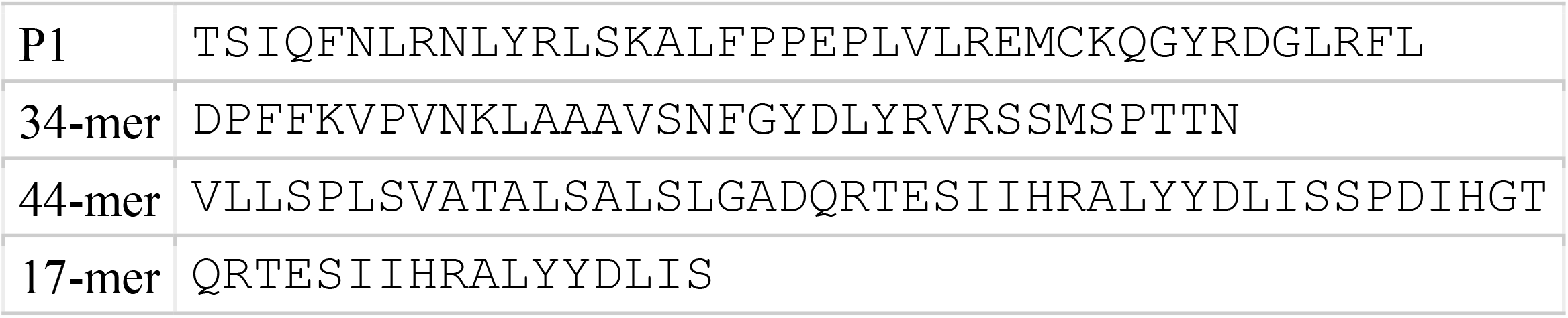
Sequences of peptides used in this study.

